# Anti-Polyamine Therapy Restrains Kidney Cyst Growth in an Orthologous Mouse Model of Autosomal Dominant Polycystic Kidney Disease

**DOI:** 10.64898/2026.06.02.729335

**Authors:** Katharina Hopp, Marie-Louise T Monaghan, Sizhao Lu, Timothy A Fields, Michael P Vitek, Katherine I Swenson-Fields

## Abstract

Autosomal Dominant Polycystic Kidney Disease (ADPKD) is the most common monogenic kidney disease worldwide. Tolvaptan is currently the only approved intervention to slow kidney cyst growth, but its indication is limited to rapidly progressing ADPKD and use compromises quality of life. Hence, new therapeutic options are of high clinical importance. Metabolic dysregulation is a hallmark of kidney cyst growth, including reprogramming of arginine metabolism and polyamine production. Sustained increases in polyamine synthesis drive pathological epithelial cell proliferation, tissue remodeling, and activation of M2-like macrophages, all known drivers of PKD. Difluoromethylornithine (DFMO) is an FDA approved irreversible inhibitor of ornithine decarboxylase (ODC1), the rate limiting enzyme in the arginine-polyamine pathway. Here, we treated C57Bl6/J p.R3277C (*Pkd1*^RC/RC^) mice for five months with DFMO-supplemented drinking water to test if DFMO treatment can slow kidney cyst growth in an orthologous model of ADPKD that mimics the disease pathophysiology seen in patients. At study end, *Pkd1*^RC/RC^ mice treated with DFMO presented with significantly reduced percent kidney weight normalized to body weight as well as kidney cystic index and cyst number compared to control. Kidney RNA-seq analyses revealed correction of many genes and pathways perturbed in PKD post DFMO treatment, as well as overall heathier kidney cellular architecture as inferred by RNAseq-base cell type deconvolution. Our data highlight significant potential to repurpose DFMO for the treatment of patients with ADPKD and investigate the role of polyamines in modulating epithelial as well as myeloid cell fate in the setting of PKD.

## Text

Autosomal Dominant Polycystic Kidney Disease (ADPKD) affects ∼1 in 1,000 live births, is predominantly caused by mutations to *PKD1*/*PKD2*, and leads to kidney failure mid-life (https://pkdcure.org/). Current treatments to slow disease progression are limited to tolvaptan with a use indication for patients with rapidly progressive disease only and side effects that impact quality of life. Metabolic dysregulation is a hallmark of kidney cyst growth, including reprogramming of arginine metabolism^1^. Arginine auxotrophy is characteristic of cyst epithelial- and atrophic proximal tubule cells; arginine deprivation results in decreased epithelial cell proliferation *in vitro* and *ex vivo* metanephric organ cystogenesis^2^. Arginase 1 (Arg1), a defining marker of M2-like macrophages, is induced in murine ADPKD kidney macrophages while macrophage depletion has been shown by others and us to slow PKD in murine models^3, 4^. Arg1^+^ macrophage depletion and ARG1 activity inhibition retard murine kidney cyst growth^3^. Arginase-mediated conversion of arginine to ornithine is the first step in the arginine-to-polyamine pathway. Next, ornithine decarboxylase (ODC1) catalyzes ornithine to putrescine. While polyamines are required for kidney epithelial and immune cell homeostasis, sustained increases in polyamine synthesis drive pathological epithelial cell proliferation, tissue remodeling, and activation of M2-like macrophages in kidney disease^5^. Of note, we have reported increased arginine–polyamine pathway activity and polyamine production in *Pkd1*^RC/RC^ mice, an orthologous model of ADPKD^6^.

Difluoromethylornithine (DFMO) is an irreversible inhibitor of ODC1. DFMO is FDA-approved for high-risk neuroblastoma, a rare pediatric cancer. DFMO has a favorable safety and tolerability profile long-term, and we found that treatment with DFMO reduces cyst size in organoid cultures from ADPKD patient biopsies (WO2020/131680A1). To determine whether anti-polyamine therapy has therapeutic potential to slow PKD progression *in vivo* we treated C57Bl6/J *Pkd1*^RC/RC^ with DFMO.

C57Bl6/J *Pkd1*^RC/RC^ mice were treated with DFMO in the drinking water (665 mg/l = 133 mg/kg) from postnatal day (P) 29 to six months of age. Control (Cntl.) animal received water without drug; both groups had *ad libitum* water access. Long-term DFMO treatment did not affect body weight compared to control (end of study [mean±SEM]; Males: Cntl: 29.73g±1.87g, DFMO:29.70g±1.55g, p=0.9048 Females: Cntl: 24.98g±0.64g, DFMO:23.21g±0.29g, p=0.0656) but showed significant therapeutic efficacy in slowing PKD (**Figure 1**). We observed a significant reduction in our primary outcomes measure, percent kidney weight/body weight (%KW/BW; **Figure 1B**). Modeling the relationship between DFMO response and sex highlighted sex as a non-significant contributor to treatment response (p=0.6890), although treated females presented with greater therapeutic efficacy (%KW/BW: Cntl. vs. DFMO; Males: 9.85% decrease, p=0.1906; Females: 33.43% decrease, p=0.0079). In addition, DFMO significantly reduced kidney cystic index, average cyst size, and cyst number (**Figure 1C**) with more pronounced decreases in females vs. males (Cntl. vs. DFMO; cystic index: Males: 50.59%, p=0.0635; Females: 65.49%, p=0.0079; average cyst size: Males: 36.30%, p=0.0635; Females: 43.90%, p=0.0079; cyst number: Males: 22.40%, p=0.2143 Females: 57.90%, p=0.0079). We did not observe a significant change in blood urea nitrogen (BUN) levels comparing DFMO-treated to Cntl. *Pkd1*^RC/RC^ mice (**Figure 1D**). However, BUN levels in C57Bl6/J *Pkd1*^RC/RC^ mice were not significantly elevated compared to age/sex-mated wildtype mice (N=6; [Mean±SEM]); Wildtype: 31.70±2.22, *Pkd1*^RC/RC^ (Cntl.): 34.69±1.93, p=0.2888) indicative of no significant ADPKD-related kidney function declines at six months of age.

**Figure 1.**
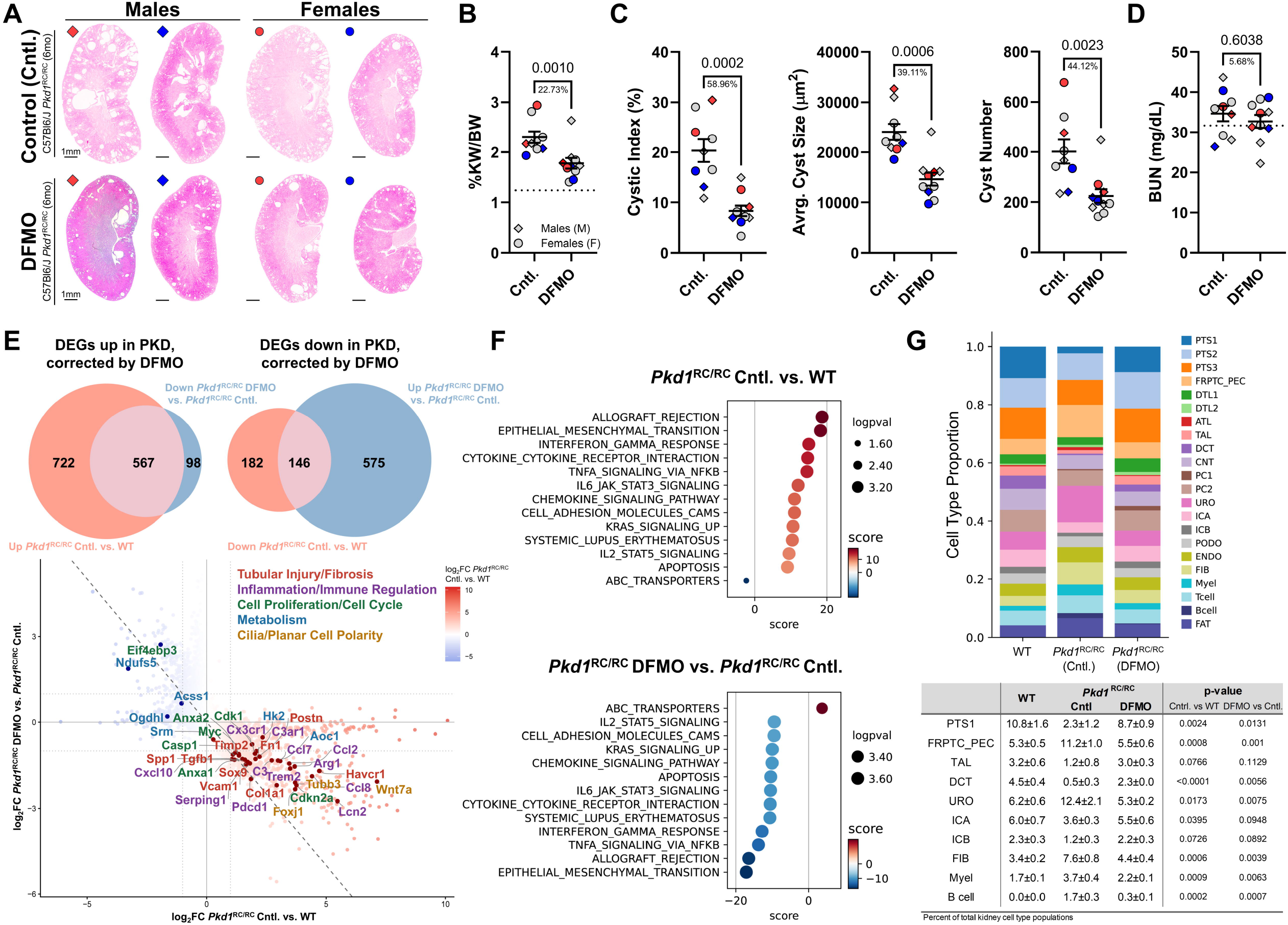
DFMO treatment slows kidney cyst growth in an orthologous mouse model of ADPKD and is associated with correction of known pathomechanistic pathways driving PKD. **(A)** Representative H&E-stained coronal kidney sections of male and female C57Bl6/J *Pkd1*^RC/RC^ mice receiving water without (Cntl.) or with DFMO. The representative animals shown in (A) are identified as colored data points in B-D, red data points = animals presenting with more severe PKD in either group or blue data points representing animals with less severe PKD in either group. Quantification of **(B)** %kidney weight/body weight (%KW/BW), **(C)** cystic index, average cyst size, and cyst number comparing animals on Cntl. or DFMO treatment. **(D)** Blood urea nitrogen (BUN) levels of animals in the Cntl. or DFMO treatment group. All parameters were collected at end of study. Each datapoint represents a single experimental animal and mean ± SEM is shown. N = 9-10 animals/group; dotted line represents the average %KW/BW or BUN of N = 3 male and 3 female C57Bl6/J wildtype age-matched mice as a means of a physiological baseline measurement. Statistics: non-parametric Mann-Whitney test, p-values are shown. Percent decrease comparing the means of Cntl vs. DFMO are shown below the p-values. | Bulk RNAseq analyses of kidney tissue harvested from WT, *Pkd1*^RC/RC^ Cntl., and *Pkd1*^RC/RC^ DFMO mice (N = 4 per group; 2 males, 2 females). **(E)** Venn diagrams showing differentially expressed genes (DEGs) of pairwise comparisons (*Pkd1*^RC/RC^ Cntl. vs. WT and *Pkd1*^RC/RC^ DFMO vs. *Pkd1*^RC/RC^ Cntl.), highlighting DEGs that were up or down in the setting of PKD and corrected by DFMO treatment. Scatter plot showing log2 fold-change (log2FC) values for all genes with significant differential expression in at least one contrast: *Pkd1*^RC/RC^ Cntl. vs. WT (x-axis) and *Pkd1*^RC/RC^ DFMO vs. *Pkd1*^RC/RC^ Cntl. (y-axis) for all DEGs that were corrected (713 DEGs); selected genes that have been published to be linked to PKD pathogenesis are highlighted. **(F)** Dotplots show the top perturbed pathways in *Pkd1*^RC/RC^ Cntl. vs. WT and top downregulated pathways in *Pkd1*^RC/RC^ DFMO vs. *Pkd1*^RC/RC^ Cntl. filtered for pathways containing DEGs that were dysregulated in the setting of PKD and corrected by DFMO and showing opposite directional enrichment between the two contrasts. **(G)** Cell-type deconvolution of bulk RNA-seq data was performed using CIBERSORTx with a published mouse ADPKD kidney snRNA-seq dataset (GSE268494) as reference. Stacked bar plots show predicted cell-type proportions per condition. Statistical comparisons are shown for select cell types in the table below. Abbreviations: PTS1/S2/S3, proximal tubule segments 1/2/3; FRPTC_PEC, failed repair proximal tubular cells/parietal epithelial cells; DTL1/DTL2/ATL, descending thin limbs 1/2 and ascending thin limb of Henle’s loop; TAL, thick ascending limb of Henle’s loop; DCT, distal convoluted tubule; CNT, connecting tubule; PC1/2, principal cells 1/2; URO, uroepithelial cells; ICA/ICB, Type A/B intercalated cells; PODO, podocytes; ENDO, endothelial cells; FIB, fibroblasts; Myel, myeloid cells; FAT, adipocytes.

To understand mechanisms driving therapeutic efficacy of DFMO in slowing PKD, we performed RNAseq of age-matched/end-of-study C57Bl6/J wildtype, *Pkd1*^RC/RC^ Cntl., and DFMO-treated kidneys (N=2M/2F/group/genotype). We identified a total of 567 differentially expressed genes (DEGs) that were upregulated in the setting of PKD (*Pkd1*^RC/RC^ Cntl. vs. WT) and downregulated by DFMO (*Pkd1*^RC/RC^ DFMO vs. *Pkd1*^RC/RC^ Cntl.); 146 DEGs were downregulated in the setting of PKD and upregulated by DFMO (**Figure 1E**). Many of the corrected genes are known to modulate PKD pathogenesis. Pathway analyses comparing *Pkd1*^RC/RC^ Cntl. vs. WT and *Pkd1*^RC/RC^ DFMO vs. *Pkd1*^RC/RC^ Cntl. with a focus on the 713 DEGs highlight correction of multiple pathways known to drive PKD progression and/or regulated by polyamines (e.g., inflammation, epithelial-mesenchymal transition, apoptosis; **Figure 1F**). To understand how kidney cellular architecture was impacted by DFMO treatment we performed cell type deconvolution using CIBERSORTx (**Figure 1G**). Interestingly, post DFMO treatment, *Pkd1*^RC/RC^ kidneys were comprised of significantly reduced numbers of failed-repair proximal tubule cells, fibroblasts, and myeloid cells, all linked to PKD progression. Further, correlative with a healthier kidney, DFMO-treated *Pkd1*^RC/RC^ kidneys had a significant increase in proximal tubule segment 1 and distal convoluted tubule cells. B cell numbers were also significantly decreased post DFMO treatment, but their role in PKD remains unexplored.

Together, our data highlight significant potential to repurpose DFMO for the treatment of patients with ADPKD. Notably, it outperformed tolvaptan in clinical efficacy as reported in a comparable preclinical study using the same animal model of ADPKD (% decrease in %KW/BW; Tolvaptan: 9.63%, DFMO: 22.73%)^7^. Further investigations are warranted into the role of polyamines in modulating epithelial as well as myeloid cell fate in the setting of PKD.

## Disclosure

KH receives royalties for industry use of the *Pkd1*^RC/RC^ mouse model in concordance with Mayo Clinic Ventures regulations (Mayo Technology Case 2012-14). MPV is employed by Regennova, Inc.

## Funding

Support was provided by the Zell Family Foundation (KH), the PKD Foundation Fellowship grant 1282646 (MLM), NIH NIDDK R41DK125183 and R44DK141418 (MPV, TAF, KISF and KH).

## Acknowledgement

The *Pkd1*^RC/RC^ mouse model was provided by Peter C. Harris, Mayo Clinic Robert M. and Billie Kelley Pirnie Translational Polycystic Kidney Disease Research Center.

## Author Contributions

Conceptualization: KH, TAF, MPV, KISF

Data Curation: KH, MTM, LS, MPV, KISF

Investigation: KH, MTM, LS, MPV, KISF

Methodology: MTM, KS, MPV, KISF

Writing: KH, MTM, MPV

## Data Sharing Agreement

All RNAseq data is publicly available; deposited at GEO (GSE333193). Code used for analyses can be found at github (https://github.com/SizhaoLu/RC_DFMO_analysis_2026). Tables outlining top differentially expressed genes/pathways are uploaded to zenodo (https://doi.org/10.5281/zenodo.20128142).

## Supplemental Digital Content

Supplemental Methods.

## Notes

### Competing Interest Statement

KH receives royalties for industry use of the Pkd1RC/RC mouse model in concordance with Mayo Clinic Ventures regulations (Mayo Technology Case 2012-14). MPV is employed by Regennova, Inc.

## References

1. Nowak KL, Hopp K: Metabolic Reprogramming in Autosomal Dominant Polycystic Kidney Disease: Evidence and Therapeutic Potential. Clin J Am Soc Nephrol, 15: 577–584, 2020 10.2215/CJN.13291019

2. Trott JF, Hwang VJ, Ishimaru T, Chmiel KJ, Zhou JX, Shim K, et al.: Arginine reprogramming in ADPKD results in arginine-dependent cystogenesis. Am J Physiol Renal Physiol, 315: F1855–F1868, 2018 10.1152/ajprenal.00025.2018

3. Yang Y, Chen M, Zhou J, Lv J, Song S, Fu L, et al.: Interactions between Macrophages and Cyst-Lining Epithelial Cells Promote Kidney Cyst Growth in Pkd1-Deficient Mice. J Am Soc Nephrol, 29: 2310–2325, 2018 10.1681/ASN.2018010074

4. Zimmerman KA, Hopp K, Mrug M: Role of chemokines, innate and adaptive immunity. Cell Signal, 73: 109647, 2020 10.1016/j.cellsig.2020.109647

5. Luo D, Lu X, Li Y, Xu Y, Zhou Y, Mao H: Metabolism of Polyamines and Kidney Disease: A Promising Therapeutic Target. Kidney Dis (Basel), 9: 469–484, 2023 10.1159/000533296

6. Hopp K, Kleczko EK, Gitomer BY, Chonchol M, Klawitter J, Christians U, et al.: Metabolic reprogramming in a slowly developing orthologous model of polycystic kidney disease. Am J Physiol Renal Physiol, 322: F258–F267, 2022 10.1152/ajprenal.00262.2021

7. Hopp K, Hommerding CJ, Wang X, Ye H, Harris PC, Torres VE: Tolvaptan plus pasireotide shows enhanced efficacy in a PKD1 model. J Am Soc Nephrol, 26: 39–47, 2015 10.1681/ASN.2013121312

