## Supplemental Methods for "Anti-Polyamine Therapy Restrains Kidney Cyst Growth in an Orthologous Mouse Model of Autosomal Dominant Polycystic Kidney Disease"

### Supplemental Method

*Ethical Statement/Murine Housing*

All animal procedures were performed in an AAALAC-accredited facility in accordance with the Guide for the Care and Use of Laboratory Animals and approved by the University of Kansas Medical Center Institutional Animal Care and Use Committee and the NIH Office of Laboratory Animal Welfare, Division of Assurances under identification number A8585-04. From birth until postnatal day (P) 21 pups were co-housed with their parents. At P21 mice were weaned. Experimental animals were group-housed, separated by sex, with a maximum of five animals per cage. Animals were maintained on a standard diet (Teklad Global 18% Protein Rodent). Food plus water were freely available; see Treatment for details on water composition. All cages were sterile; the vivarium maintained a temperature of ~72ºF, humidity of ~37%, and light cycle of 8pm off, 6am on.

C57Bl6/J *Pkd1*$^{RC/RC}$ mice were obtained from the Mayo Clinic (Dr. Peter C. Harris) via an approved MTA. This genetic model contains a hypomorphic mutation located within the *Pkd1* loci and is widely accepted as an immunocompetent model of ADPKD. The genotypes of all experimental animals were PCR confirmed using DNA extracted from tail clips. The *Pkd1* p.R3277C genotyping protocol was performed as previously published[1].

*Study Design*

The study protocol was established prior to study initiation. C57Bl6/J *Pkd1*$^{RC/RC}$ mice were assigned to experimental groups (Control [Cntl.] or DFMO) at age of weaning (P21); littermates were separated by experimental group. Randomization to study groups was based on body weight at time of weaning assuring comparable mean ± standard error of the mean (SEM) between groups. The only exclusion criteria was a weaning weight of <10g of body weight. An experimental unit (number; N) was defined as a single animal independent of sex. The study design was based on a proof-of-concept design; hence no formal power calculation to determine sample size was executed prior study initiation. Treatment was initiated at P29 and animals were euthanized for outcome measures at six months of age.

*Treatment*

Treatment started at P29 and continued till the animals reached 6 months of age. Experimental animals in the control group (Cntl.) received hyperchlorinated reverse osmosis water *ad libitum*. Experimental animals in the DFMO group received the same water supplemented with 665 mg/l DFMO (CAS 96020-91-6, Ambeed Inc., Arlington Hts, IL) to achieve a continuous dose of 133 mg/kg DFMO *ad libitum*. Animals were monitored weekly for physical appearance to assess potential unexpected events associated with the treatment regimen or progression of PKD. Humane endpoints to reduce pain and suffering of the animals were determined to be inability to eat/drink or physical appearance changes (hunched posture, ruffled fur, or severe dehydration) as well as behavior changes (severe lethargy, lack of grooming, or inability to ambulate) as predetermined prior study start. No unexpected events requiring euthanasia based on the set humane endpoints occurred during the study design.

*Murine Tissue Harvest/Processing*

Experimental animals were weighed, euthanized by isoflurane exposure overdose followed by opening of the chest cavity. Blood was collected via cardiac puncture and incubated on ice for 15min prior to serum isolation via centrifugation (2,500xg, 15min). The kidneys were harvested and weighed. The right kidney was cut through the mid-sagittal plane, fixed in 4% paraformaldehyde, and embedded in paraffin for histological analysis. The left kidney was stored in liquid nitrogen for potential follow-up analyses i.e., RNAseq.

*Histological Analyses*

The embedded kidneys were sectioned from the mid-sagittal plane at 4μm thickness and H&E stained. Kidney image acquisition and analysis was performed as previously described[2]. Cystic index and cyst size and number were analyzed using a custom-built NIS-Elements AR v4.6 macro (Nikon). An average of the two coronal sections was established per animal and reported as an N=1.

*Blood Urea Nitrogen Measurements*

Blood Urea Nitrogen (BUN) was measured using the QuantiChrom Urea Assay Kit (BioAssay Systems, # 501078333) according to the manufacturer's protocol. A total of 5 µL of serum was analyzed per sample and all animals were analyzed in duplicates.

*Outcome measure*

All (histo-)pathological outcome measures were predefined at time of study design. The primary outcome to determine treatment efficacy to slow PKD progression in the given study design was set to be kidney weight normalized to body weight (%KW/BW). All other outcome measures indicative of treatment response (cystic index, average cyst size/number, BUN) were deemed exploratory. All data was analyzed blinded to treatment group

*RNA-seq and bioinformatics analysis pipeline*

RNA was isolated from kidney tissue flash frozen in liquid nitrogen using the Trizol method and purified as described[3]. The quality of each RNA sample was determined with an Agilent Tape Station 4200 by the University of Kansas Medical Center Genome Sequencing Facility. Poly(A)-selected RNA libraries were constructed and sequenced in paired-end mode (2 × 150 bp) by Novogene (Sacramento, CA) on an Illumina platform. Raw sequencing reads were preprocessed to trim adapter sequences and remove low-quality reads, defined as those containing ambiguous bases (N) exceeding 10% of read length or a Phred quality score (Q-score) ≤ 5 in more than 50% of bases. Filtered reads achieved a Q30 ≥ 93% across all samples and were aligned to the mouse reference genome (mm10) using STAR (v2.5.3a)[4], with a mean uniquely mapping rate exceeding 85%. Gene-level read counts, with lowly expressed genes filtered out (total counts < 10 across all samples), were used as input for differential gene expression (DGE) analysis, contrasting *Pkd1*$^{RC/RC}$ Cntl. vs. WT and *Pkd1*$^{RC/RC}$ DFMO vs. *Pkd1*$^{RC/RC}$ Cntl. with PyDESeq2 (v0.5.2)[5]. Log fold-change (logFC) shrinkage was applied prior to visualization. Differentially Expressed Genes (DEGs) corrected by DFMO were defined as those differentially expressed in opposite directions between the two contrasts (log2FC >1 or <−1, padj<0.05 in both comparisons). Pathway enrichment analysis was performed using decoupleR (v2.1.6)[6] with KEGG pathway and hallmark gene sets obtained from the Molecular Signatures Database (MSigDB). To identify top pathways corrected by DFMO, enrichment results from both contrasts were filtered for pathways that (1) contained DEGs corrected by DFMO and (2) showed opposite directionality between *Pkd1*$^{RC/RC}$ Cntl. vs. WT and *Pkd1*$^{RC/RC}$ DFMO vs. *Pkd1*$^{RC/RC}$ Cntl. (i.e., upregulated in *Pkd1*$^{RC/RC}$ Cntl. vs. WT and downregulated in *Pkd1*$^{RC/RC}$ DFMO vs. *Pkd1*$^{RC/RC}$ Cntl., or vice versa). Cell-type deconvolution of bulk RNA-seq data was performed using CIBERSORTx[7] (https://cibersortx.stanford.edu). A published, annotated mouse PKD single-cell RNA-seq dataset (GSE268494) was used as the reference. To construct the signature matrix, the reference dataset was downsampled to a maximum of 200 cells per cell type, and analysis was restricted to genes present in both the single-cell reference and bulk RNA-seq datasets. Cell-type fraction estimation was performed with S-mode batch correction enabled and 100 permutations.

*Data availability*

All data supporting the findings of this study are publicly available. Raw sequencing data generated in this study have been deposited in the NCBI Gene Expression Omnibus (GEO) under accession number GSE333193 (https://www.ncbi.nlm.nih.gov/geo/query/acc.cgi?acc=GSE333193). Code used for analyses can be found at github (https://github.com/SizhaoLu/RC_DFMO_analysis_2026). Processed data used to generate the figures in this manuscript are available through the Zenodo repository (https://doi.org/10.5281/zenodo.20128142).

*Statistical Analyses*

All analyses were performed using JMP Pro 18.2.2(SAS) or PRISM 11.0.1 (Graphpad Software). Data are presented as the mean ± standard error of the mean (SEM); single data points are depicted. Pairwise comparisons were made using a non-parametric Mann-Whitney test. The effect of sex as an outcome-driving variable was determined using JMP "Fit Model" a formula-based modeling using Standard Least Squares (regression and ANOVA) based on %KW/BW. Comparisons between WT, *Pkd1*$^{RC/RC}$ Cntl., and *Pkd1*$^{RC/RC}$ DFMO CIBERSORTx[7] outputs were analyzed using One-way ANVOA with Tukey post-hoc test to correct for multiple comparisons. All p-values are denoted on the figure.

**Supplemental References**


1. Hopp K, Ward CJ, Hommerding CJ, Nasr SH, Tuan HF, Gainullin VG, et al.: Functional polycystin-1 dosage governs autosomal dominant polycystic kidney disease severity. *J Clin Invest,* 122**:** 4257-4273, 2012 10.1172/JCI64313
2. Kleczko EK, Marsh KH, Tyler LC, Furgeson SB, Bullock BL, Altmann CJ, et al.: CD8(+) T cells modulate autosomal dominant polycystic kidney disease progression. *Kidney Int,* 94**:** 1127-1140, 2018 10.1016/j.kint.2018.06.025
3. Swenson-Fields KI, Ward CJ, Lopez ME, Fross S, Heimes Dillon AL, Meisenheimer JD, et al.: Caspase-1 and the inflammasome promote polycystic kidney disease progression. *Front Mol Biosci,* 9**:** 971219, 2022 10.3389/fmolb.2022.971219
4. Dobin A, Davis CA, Schlesinger F, Drenkow J, Zaleski C, Jha S, et al.: STAR: ultrafast universal RNA-seq aligner. *Bioinformatics,* 29**:** 15-21, 2013 10.1093/bioinformatics/bts635
5. Muzellec B, Telenczuk M, Cabeli V, Andreux M: PyDESeq2: a python package for bulk RNA-seq differential expression analysis. *Bioinformatics,* 39, 2023 10.1093/bioinformatics/btad547
6. Badia IMP, Velez Santiago J, Braunger J, Geiss C, Dimitrov D, Muller-Dott S, et al.: decoupleR: ensemble of computational methods to infer biological activities from omics data. *Bioinform Adv,* 2**:** vbac016, 2022 10.1093/bioadv/vbac016
7. Newman AM, Steen CB, Liu CL, Gentles AJ, Chaudhuri AA, Scherer F, et al.: Determining cell type abundance and expression from bulk tissues with digital cytometry. *Nat Biotechnol,* 37**:** 773-782, 2019 10.1038/s41587-019-0114-2
